# Cognitive impairment and metabolite profile alterations in the hippocampus and cortex of male and female mice exposed to a fat and sugar-rich diet are normalized by diet reversal

**DOI:** 10.1101/2021.04.27.441603

**Authors:** Alba M. Garcia-Serrano, Adélaïde A. Mohr, Peter Spégel, João M.N. Duarte

## Abstract

Diabetes impacts on brain metabolism, structure and function. Alterations in brain metabolism have been observed in obesity and diabetes models induced by exposure to diets rich in saturated fat and/or sugar, and have been linked to memory impairment. However, it remains to be determined whether brain dysfunction induced by obesogenic diets results from permanent brain alterations. We tested the hypothesis that an obesogenic diet (high-fat and high-sucrose diet; HFHSD) causes reversible changes in hippocampus and cortex metabolism and alterations in behavior. To test this hypothesis, mice were exposed to HFHSD for 24 weeks or for 16 weeks followed by 8 weeks of diet normalization. Development of the metabolic syndrome, changes in behavior, and brain metabolite profiles by ^1^H magnetic resonance spectroscopy (MRS) were assessed longitudinally. Control mice were fed an ingredient-matched low-fat and low-sugar diet. Mice fed a HFHSD developed obesity, glucose intolerance and insulin resistance, with a more severe phenotype in male than female mice. Relative to controls, both male and female HFHSD-fed mice showed increased anxiety-like behavior, impaired memory in object recognition tasks, but preserved working spatial memory as evaluated by spontaneous alternation in a Y-maze. Alterations in the metabolite profiles were observed both in the hippocampus and cortex, but were more distinct in the former. HFHSD-induced metabolic changes included altered levels of lactate, glutamate, GABA, glutathione, taurine, *N*-acetylaspartate, creatine and choline. Notably, HFHSD-induced metabolic syndrome, anxiety, memory impairment, and brain metabolic alterations recovered upon diet normalization for 8 weeks. We conclude that cortical and hippocampal derangements induced by long-term HFHSD consumption are reversible rather than being the result of permanent tissue damage.

## Introduction

The incidence of neurodegenerative diseases is increasing worldwide in its sporadic forms, which is likely due to increased longevity and unhealthy lifestyles (Popa-Wagner *et al.*, 2020). In particular, obesity is associated closely to comorbidities such as cardiovascular disease, metabolic syndrome and type 2 diabetes (T2D) (Askari, 2020), disorders that impact the brain and have been linked to mild cognitive impairment, Alzheimer’s disease or vascular dementia (Frisardi *et al.*, 2010; Spauwen *et al.*, 2013; de la Monte, 2017). Mechanisms of brain dysfunction linked to obesity and metabolic disorders are suggested to include glucose toxicity, vascular dysfunction, mitochondrial damage, oxidative stress, synaptic failure or neuroinflammation (Garcia-Serrano & Duarte, 2020). In rodent models, obesity and metabolic syndrome have been linked with alterations in brain function (de Bem *et al.*, 2021) and metabolism (Garcia-Serrano & Duarte, 2020).

Proton magnetic resonance spectroscopy (MRS) enables non-invasive profiling of regional metabolic profiles (Duarte *et al.*, 2012). Diabetes and associated comorbidities have been shown to cause variations in brain concentrations of *N*-acetylaspartate (NAA), creatine, choline, *myo*-inositol, glutamate and glutamine (Duarte *et al.*, 2016; Zhao *et al.*, 2018). Alterations in metabolism have also been observed in multiple diabetes models (van der Graaf *et al.*, 2004; Duarte *et al.*, 2009, 2019; Girault *et al.*, 2019; Wang *et al.*, 2012), and in rodents that develop metabolic syndrome upon long-term exposure to obesogenic diets (Ribeiro *et al.*, 2018; Lizarbe, Soares *et al.*, 2019; Lizarbe, Cherix *et al.*, 2019). While the metabolic syndrome caused by long-term exposure to obesogenic diets is reversible (Soares *et al.*, 2018), it remains to be determined whether associated alterations in brain metabolism are plastic, or the result from major structural injury to brain tissue. We aimed at testing the hypothesis that a long-term high-fat and high-sucrose diet (HFHSD) exposure causes reversible alterations of brain function (behavior) and metabolism.

## Materials and Methods

### Animals

Experiments were performed according to EU Directive 2010/63/EU under approval of the Malmö/Lund Committee for Animal Experiment Ethics (permit number 994/2018), and are reported following the ARRIVE guidelines (Animal Research: Reporting In Vivo Experiments, NC3Rs initiative, UK). The sample size was estimated from previous MRS studies (Gapp *et al.* 2017; Lizarbe, Soares *et al.*, 2019).

Male and female C57BL/6J mice (8-weeks old) were purchased from Taconic Biosciences (Köln, Germany). Mice were housed in groups of 4-5 animals on a 12h light-dark cycle with lights on at 07:00, room temperature of 21-23 °C and humidity at 55-60%. Mice were habituated to the facility for 1 week upon arrival.

Mice were randomly assigned to 3 experimental groups, receiving either a control diet (CD, 10%-fat diet, n=12 males + 15 females), a high-fat and high-sucrose diet (HFHSD, n=13 males + 13 females) consisting of 60%-fat diet feeding plus access to a 20%(w/v) sucrose in drinking water from 10 weeks of age and during 6 months, or a reversed diet (reversed, n=9 males + 10 females) that consisted of HFHSD feeding for 4 months followed by CD feeding for 2 months. Mice receiving HFHSD also had access to sugar-free water. Food and water were provided *ad libitum.*

This study used open-source diets from Research Diets (New Brunswick, NJ-USA): a lard-based diet with 60% kcal of fat (D12492) and a control diet containing 10% kcal of fat (D12450J), with total energy of 5.21 and 3.82 kcal/g, respectively. Both diets contain 20% kcal from protein and 7% kcal from sucrose, and the remaining calories are from carbohydrates (maltodextrin and starch).

### Magnetic Resonance Spectroscopy (MRS)

MRS was performed at baseline and at 1, 2, 4, 8, 16 and 24 weeks after initiation of the diet. For 6 mice in each group, MRS was only conducted at 16 and 24 weeks. All experiments were performed on a preclinical 9.4T Bruker BioSpec AV III (Bruker, Ettlingen, Germany) with an effective bore size of 86 mm and gradient strength of 670mT/m, using ParaVision 6.0.1 (RRID:SCR_001964), and equipped with a ^1^H quadrature transmit/receive cryoprobe. Anaesthesia was induced with 3% isoflurane (Vetflurane, Virbac, Carros, France) in a 1:1 (v/v) O_2_:N_2_O gas mixture. Then mice were positioned onto a MRS-compatible bed with a teeth bar and two ear bars for stereotaxic head fixation. Anesthesia was delivered through a home-build mask at variable rate of 1-2% isoflurane for maintaining respiration at 60-90 breaths per minute. Body temperature was kept between 36-37°C by means of warm water circulation. Breathing rate and body temperature were continuously recorded with the SA Instruments (Stony Brook, NY, USA) monitoring system.

After ensuring good head position in the scanner by means of a low resolution localizer image, T2-weighted images over the whole brain were acquired for anatomical reference using the Rapid Imaging with Refocused Echoes (RARE) sequence with repetition time TR=3.5 s, echo time TE=33 ms, 32 slices, thickness of 0.5 mm, 320×320 voxels, and FOV=14×14 mm^2^. After MAPSHIM (Bruker) and iterative linear shimming, proton MRS was acquired using STimulated Echo Acquisition Mode (STEAM) with TR=4 s, TE=3 ms, mixing time TM=20 ms, and a spectral width of 4401.41 Hz. Volumes of interest (VOI) for MRS were placed in dorsal hippocampus (1.8 mm x 1.2 mm x 1.5 mm) and cortex (4 mm x 0.8 mm x 1.5 mm).

Water-suppressed spectra were acquired in 20 and 12 blocks of 16 scans in the hippocampus and cortex, respectively. An unsuppressed water spectrum from the same VOI was acquired in one block of 16 averages. After block alignment and summation in MATLAB (MathWorks, Natick, MA), metabolite concentrations were determined with LCModel v.6.3-1A (Stephen Provencher Inc., Oakville, Ontario-Canada; RRID:SCR_014455), including a macromolecule spectrum in the database and using the unsuppressed water signal measured from the same VOI as internal reference (Duarte *et al.*, 2014). The following metabolites were included in the LCModel analysis: alanine, ascorbate, aspartate, β-hydroxybutyrate, creatine, γ-aminobutyrate (GABA), glutamine, glutamate, glutathione, glycine, glycerophosphorylcholine (GPC), glucose, lactate, *myo*-inositol, *N*-acetylaspartate, *N*-acetylaspartylglutamate (NAAG), phosphorylethanolamine (PE), phosphorylcholine (PCho), phosphocreatine, *scyllo*-inositol, and taurine. After LCModel analysis, metabolites with Cramér-Rao lower bound (CRLB) larger than 30% were disregarded as they did not fulfil reliability criteria. Namely, β-hydroxybutyrate and *scyllo*-inositol were excluded, and phosphorylcholine and glycerophosphorylcholine were analyzed as total choline (PCho+GPC).

Spectra showing extra-cerebral lipid contamination were excluded from the analysis (36 out 828 MRS scans). At the end of the study, one mouse was excluded due to simultaneous increase of glutamine levels and decreased osmolytes in both brain areas and at all MRS time points, which suggests occurrence of a congenital portosystemic shunt (Cudalbu *et al.*, 2013).

### Behavior

Mice were allowed to acclimatize to the testing room for 1 hour. Tests were performed from 9:00 to 18:00, with light adjusted to an illuminance of 15 lx in each apparatus.

Spontaneous alternations were observed in a Y-maze as surrogate of working memory performance (Duarte, Agostinho *et al*, 2012), at baseline and after 1, 2, 4, 8, 16 and 24 weeks of treatment. The Y-maze arms were 30 cm x 15 cm x 5 cm (length x height x width), and converged to the center at an 120° angle. Mice were placed in one arm and allowed to freely explore the maze for 8 minutes. Complete spontaneous alternations were defined as successive entries into the three arms, and expressed relative to the total possible alternations in the test. The total number of entries was used to access locomotor activity and exploratory behavior.

Object recognition tasks were adapted from de Paula *et al*. (2020), in which memory is tested based on the spontaneous tendency of rodents familiarized with two objects to explore a novel object (novel object recognition, NOR) or an object relocated in space (novel location recognition, NLR). Exploration was recorded by an infrared camera in a cubic arena with a side length of 50 cm. Mice were first habituated to the empty arena for 8 minutes. Arena exploration was analyzed for total walk distance, number of crossings between arena quadrants and immobility time, as well as exploration of the arena center at 6 cm from the walls. Thereafter, NLR was assessed by placing the mice in the arena with two identical objects, and allowed to explore them for 5 minutes (familiarization phase). Mice were then removed from the arena for 1 hour (retention phase), and reintroduced for 5 minutes but with one object relocated to a different quadrant in the arena (recognition phase). For NOR, two new identical objects were used in the familiarization phase, and one of them was replaced by a novel object during the recognition phase. Time exploring each object was measured.

### Glucose tolerance test (GTT) and hormone analyses

A GTT was performed before (baseline) and after 1, 2, 4, 8, 16 and 24 weeks on the diets. Food was removed for 6 hours starting at 08:00. Thereafter, a blood sample was collected from the vena saphena to determine plasma hormone levels. Glycemia was measured from tail tip blood with the Breeze glucometer (Bayer, Zürich, Switzerland). Then mice were given 2 g/kg glucose i.p. from a 30%(w/v) solution in saline, followed by determination of glucose levels at 15, 30, 60, 90, and 120 minutes. Commercially available ELISA kits were used to determine plasma concentrations of insulin (#10-1247-10, Mercodia, Uppsala, Sweden; RRID: AB_2889906) and leptin (#ab100718, Abcam, Cambridge, United Kingdom; RRID: AB_2889903).

### Statistics

Data were analyzed by ANOVA followed by independent comparisons with the Fisher’s least significant difference (LSD) test in Prism 9.0.2 (GraphPad, San Diego, CA-US; RRID:SCR_002798). Principal component analysis (PCA) was conducted in R 3.6.2 using prcomp (RRID:SCR_001905). Results are presented as mean±SD unless otherwise stated.

## Results

### Food intake and metabolic phenotype

Mice on HFHSD had a higher caloric intake as compared to mice on the control diet (figure 1B), which was due increased consumption of fat and sucrose, rather than carbohydrates or protein (figure 1C). Consequently, mice fed HFHSD became overweight, but then returned to normal levels after a transition to the control diet (figure 1D). Weight gain from baseline to the end of the study was higher in HFHSD-fed mice, but not in the group of mice for which the diet was reversed, relative to controls (diet P<0.001; gender P<0.001; interaction P=0.657; figure 1E).

**Figure 1.**
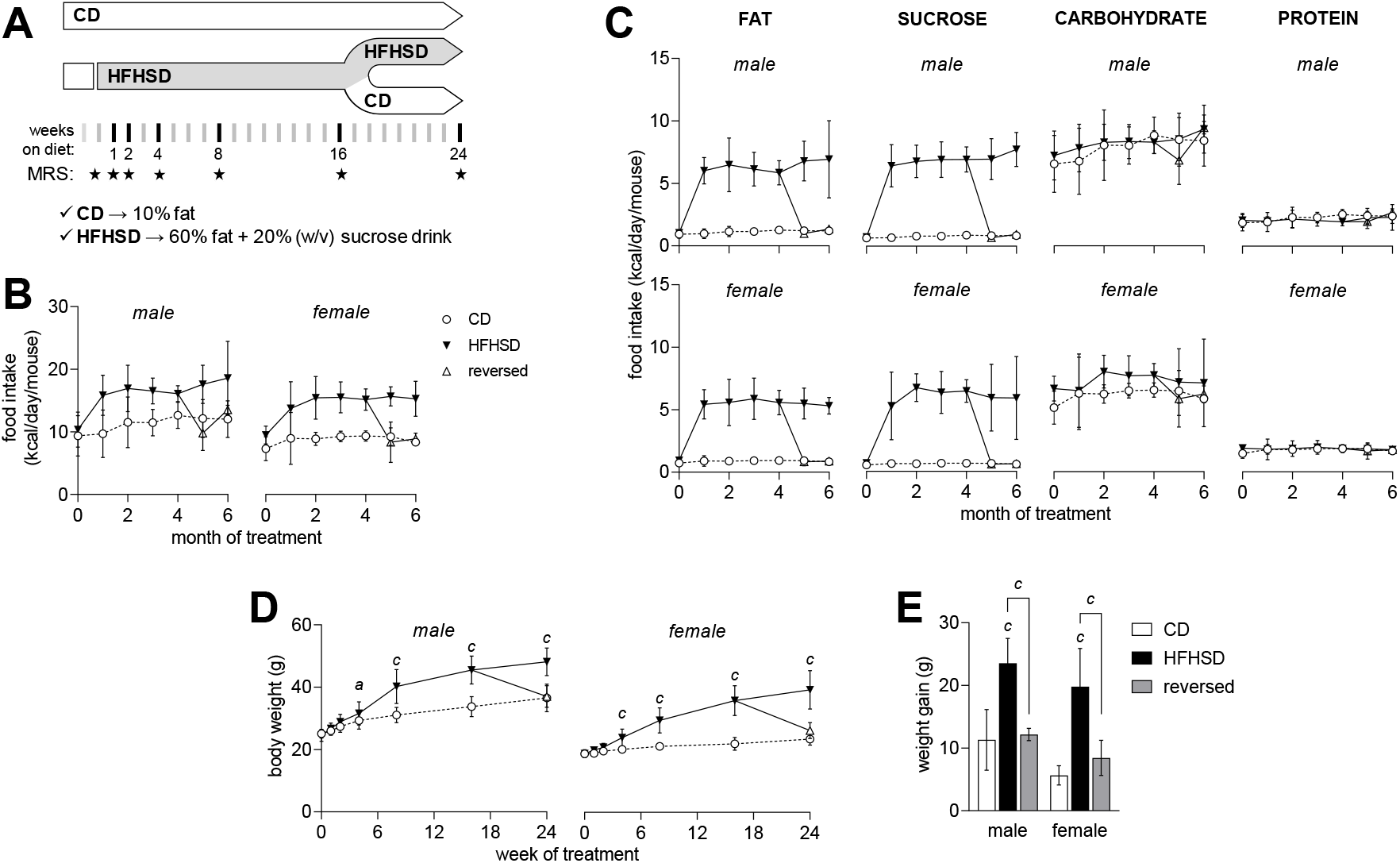
Study design and caloric intake in each month of the treatment. Mice were acclimatized under control diet (CD) at 8 weeks of age, and exposed to HFHSD for 4 or 6 months starting at 10 weeks of age (A). A group of mice had the diet reversed to control after 4 months under HFD (reversed). MRS scans took place at baseline, and then at weeks 1, 2, 4, 8, 16 and 24 of the treatment (stars in timeline of panel A). Average caloric intake from increased during HFHSD-feeding (B) due to increased fat and sucrose intake (C). Relative to CD, HFHSD feeding resulted in increased body weight (D) and an over 2-fold larger weight gain from baseline to 6 months of treatment (E), which is fully reversed by diet normalization. Data is mean±SD. Letters over data-points indicate significant differences relative to CD or as indicated (^*a*^ P<0.05, ^*b*^ P<0.01, ^*c*^ P<0.001) based on Fisher’s LSD post hoc comparison for significant effects of diet or interaction between diet and time, as assessed by ANOVA.

In line with previous studies in mice fed other obesogenic diets (Soares *et al.*, 2018), the HFHSD caused sex-dependent imbalances in glucose homeostasis, which were fully recovered after diet reversal (figure 2). Interestingly, at the end of the 24 weeks of treatment, HFHSD-associated glucose intolerance was more severe in male than female mice (figure 2B-C), and insulin resistance was negligible in females (figure 2F). Glucose clearance during the GTT (figure 2A), as defined by both the glucose area-under-the-curve (at 24 weeks: diet P<0.001; gender P<0.001; interaction P=0.056; figure 2B) and glucose levels at 2 hours post-bolus (at 24 weeks: diet P<0.001; gender P<0.001; interaction P=0.025; figure 2C), was impaired by HFHSD feeding.

**Figure 2.**
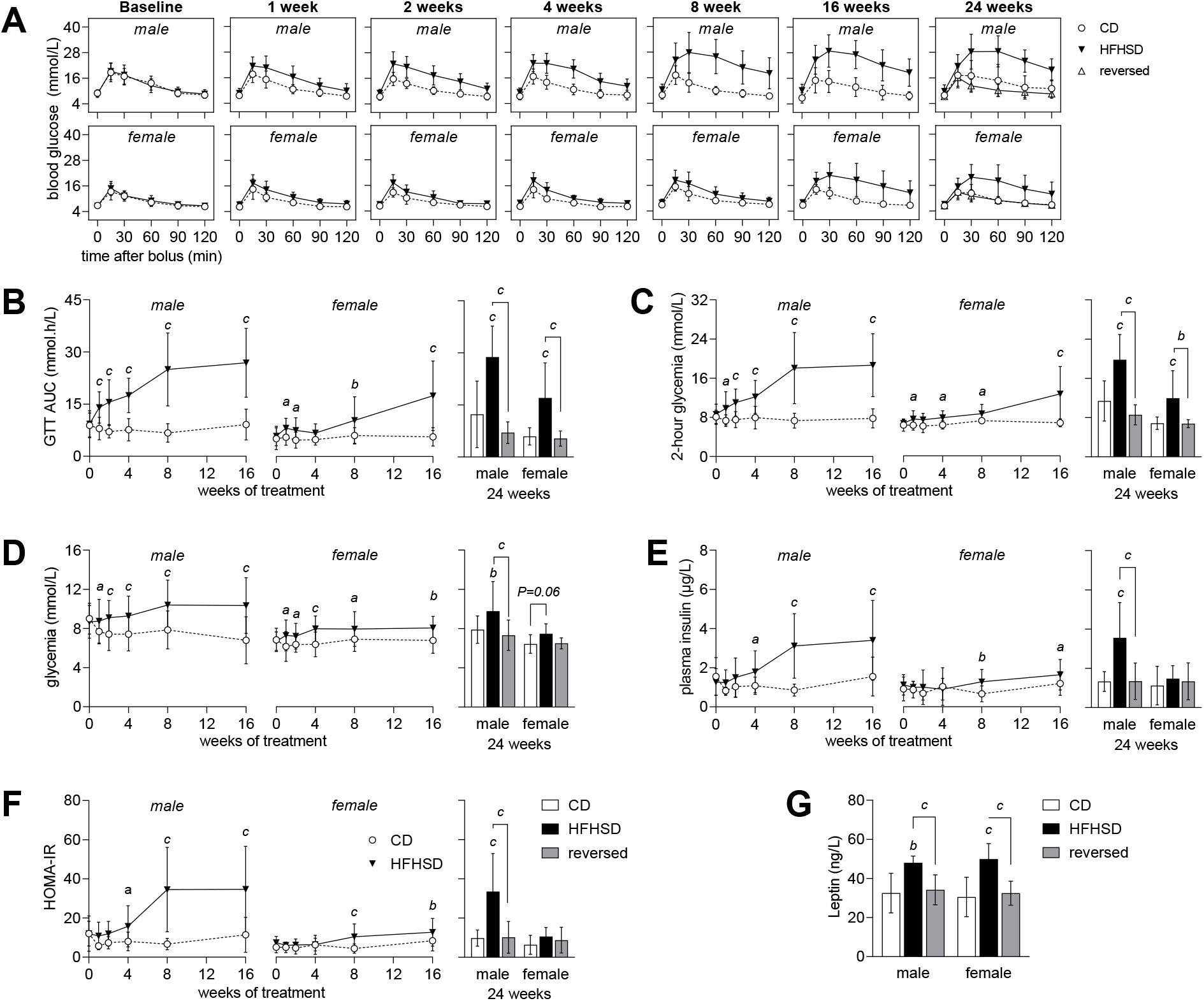
Glycemic regulation in mice from control (CD), HFHSD and reversed groups at baseline and after 1, 2, 4, 8, 16 and 24 weeks of the treatment. Glucose clearance (A) in a glucose tolerance test (GTT) was reduced by HFHSD feeding, as evidenced by both increased area under the curve (AUC) of the GTT (B) and increased glycemia 2 hours after the glucose bolus (C). Glycemia (D), plasma insulin (E) and HOMA-IR (F) after a 6-hour fasting period indicate insulin resistance in HFHSD-fed male mice. Compared to controls, plasma leptin was increased in HFHSD-fed mice and normalized by diet reversal (G). Data is mean±SD. Letters over data-points indicate significant differences relative to CD or as indicated (^*a*^ P<0.05, ^*b*^ P<0.01, ^*c*^ P<0.001) based on Fisher’s LSD post hoc comparison following presence of significant effects of diet or interaction between diet and time in ANOVA tests.

Mice fed HFHSD showed slightly higher fasting glucose (diet P<0.001; gender P<0.001; interaction P=0.292; figure 2D) and plasma insulin levels (diet P<0.001; gender P=0.011; interaction P=0.007; figure 2E) than controls. This translated into an increased HOMA-IR, indicative of insulin resistance (diet P<0.001; gender P=0.001; interaction P=0.003; figure 2F). Plasma leptin was increased in HFHSD-fed mice but not in the diet reversal group, relative to controls (diet P<0.001; gender P=0.815; interaction P=0.801; figure 1G). Altogether, these results suggest that obesity induced by HFHSD feeding causes glucose intolerance and insulin resistance, which were normalized by diet reversal.

### HFHSD-induced behavior alterations

Having established a reversible diet-dependent effect on glycemic control, we then investigated whether metabolic syndrome was associated with behavioral alterations. Spatial working memory was evaluated by measuring Y-maze spontaneous alternation during the course of phenotype development (figure 3A). Diet had no effect on the spontaneous alternation (at 24 weeks: diet P=0.353; gender P=0.034; interaction P=0.334), or exploratory behavior depicted by number of arm entries (at 24 weeks: diet P=0.638; gender P=0.116; interaction P=0.988).

**Figure 3.**
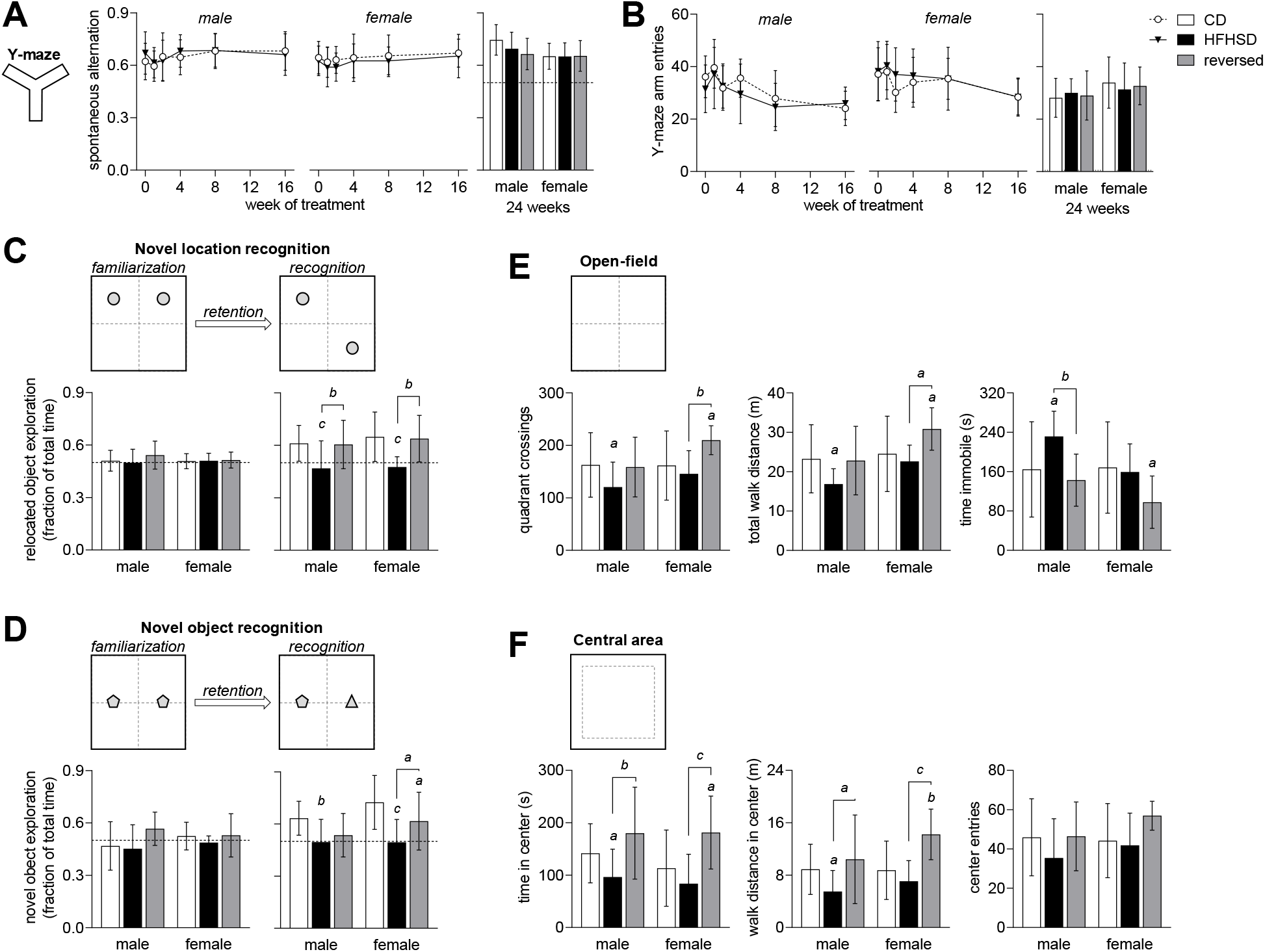
Memory impairment induced by HFHSD exposure. Spontaneous alternation (A) and number of arm entries (B) in the Y-maze were not modified during exposure to HFHSD. After familiarization with 2 objects, both novel location recognition (C) and novel object recognition (D) tasks show that mice exposed to HFHSD for 24 weeks did not display increased exploration of novel location or object. When diet was reversed, mice recovered the ability to recognize novelty in the object displacement task (C) but not in the object replacement task (D). Dashed lines in graphs represent chance (50%). Data is mean±SD. Letters over data-points indicate significant differences relative to CD or as indicated (^*a*^ P<0.05, ^*b*^ P<0.01, ^*c*^ P<0.001) based on Fisher’s LSD post hoc comparison following presence of significant effects of diet or interaction between diet and time in ANOVA tests.

Next, we examined novelty recognition at 24 weeks after introduction of the diets. After familiarization with 2 objects, mice normally tend to spend more time exploring a novel object, or an object that has been displaced in the arena. This behavior was impaired in HFHSD-fed mice when testing either novel location recognition (diet P<0.001; gender P=0.341; interaction P=0.867; figure 3C) or novel object recognition (diet P<0.001; gender P=0.059; interaction P=0.308; figure 3D). relative to HFHSD, the reversed diet group fully recovered memory performance to control levels in the novel location recognition task (figure 3C), and also improved performance in the novel object recognition task (figure 3D).

We examined open-field exploration during 8 minutes (habituation prior to object recognition tasks). HFHSD exposure reduced total distance walked in the arena (diet P=0.006; gender P=0.004; interaction P=0.224; figure 3E). Moreover, when compared to controls, male mice fed HFHSD showed reduced number of crossings across quadrants of the arena (diet P=0.008; gender P=0.047; interaction P=0.234), and increased immobility time (diet P=0.006; gender P=0.030; interaction P=0.121). Altogether, these results suggest reduced novel environment exploration upon HFHSD exposure, which is normalized after diet reversal.

Typically, rodents spend a greater amount of time exploring the periphery of the arena, next to the walls, rather than the unprotected center area (Prut & Belzung, 2003). HFHSD-fed mice spent less time (diet P<0.001; gender P=0.375; interaction P=0.716; figure 3F), and walked a shorter distance in the center of the arena (diet P<0.001; gender P=0.071; interaction P=0.256), despite similar number of center entries (diet P=0.055; gender P=0.214; interaction P=0.435). Reduced exploration of center *versus* periphery suggests increased anxiety-like behavior in HFHSD-fed mice, which recovered upon diet normalization.

### Metabolite profiles

Changes in behavior were similar in male and female mice (figure 3). Moreover, we have previously demonstrated that metabolite profiles measured by MRS in the hippocampus and cortex are similar in male and female mice (Duarte *et al.*, 2014). Thus, in order to increase the statistical power, metabolite profiles were analyzed for male and female mice grouped together. Figure 4 depicts the VOI placement in hippocampus and cortex, and respective spectra.

**Figure 4.**
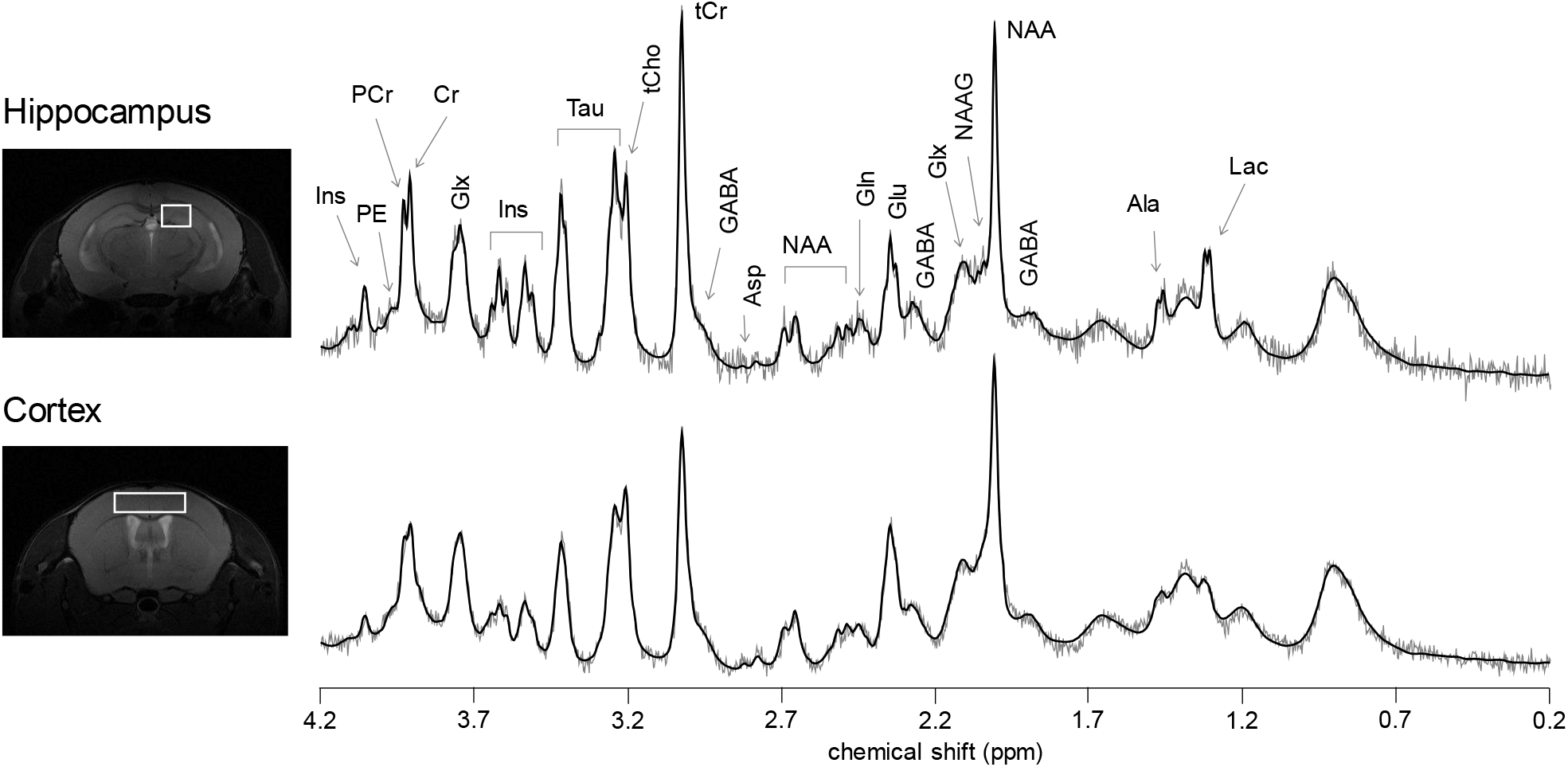
Representative location of the VOIs used for MRS in dorsal hippocampus and cortex, and respective spectra acquired with STEAM at 9.4 T (gray line) and LCModel fitting result (black line). From right to left: Ala, alanine; Lac, lactate; GABA, γ-aminobutyrate; NAA, *N*-acetylaspartate; NAAG, *N*-acetylaspartylglutamate; Glx, glutamate (Glu) + glutamine (Gln); Asp, aspartate; tCr, total creatine = creatine (Cr) + phosphocreatine (PCr); tCho, total choline = phosphorylcholine + glycerophosphorylcholine; Tau, taurine; Ins, *rnyo*-inositol; PE, phosphorylethanolamine.

HFHSD feeding altered levels of multiple metabolites in both hippocampus and cortex (table 1). In the hippocampus (figure 5), the earliest metabolic alteration triggered by HFHSD was a decrease in levels of lactate (P<0.05 *vs*. control at 4 weeks of diet) that persisted until the end of the treatment. Levels of glutamate (P<0.01), *N*-acetylaspartate (NAA; P<0.05), taurine (P<0.001) and creatine (P<0.001), increased at 8 weeks, relative the controls, and remained high throughout the intervention. At 16 weeks, concentrations of GABA (P<0.001), *myo*-inositol (P<0.01), phosphocreatine (P<0.001) and glutathione (P<0.001) increased in the HFHSD-fed mice, as compared to controls. HFHSD feeding also caused transient changes in levels of alanine at 8 weeks (P<0.01) and of *N*-acetylaspartylglutamate (NAAG) at 16 weeks (P<0.05) of treatment. All these metabolic alterations in the hippocampus were lost 8 weeks after diet reversal (table 2; figure 5).

**Table 1.** ANOVA P-values for metabolite concentrations in the hippocampus and cortex during 1 to 16 weeks of HFHSD exposure.

| Metabolite | Brain area | Interaction | Time | Diet |
| --- | --- | --- | --- | --- |
| Alanine | hippocampus | P=0.008 | P<0.001 | P=0.371 |
|  | cortex | P<0.001 | P<0.001 | P<0.001 |
| Aspartate | hippocampus | P=0.501 | P=0.764 | P=0.256 |
|  | cortex | P=0.394 | P=0.703 | P=0.572 |
| Creatine | hippocampus | P=0.061 | P<0.001 | P<0.001 |
|  | cortex | P=0.747 | P=0.029 | P=0.431 |
| Phosphocreatine | hippocampus | P<0.001 | P=0.016 | P=0.005 |
|  | cortex | P=0.743 | P<0.001 | P=0.857 |
| GABA | hippocampus | P=0.003 | P=0.200 | P=0.199 |
|  | cortex | P=0.528 | P=0.047 | P=0.880 |
| Glutamine | hippocampus | P=0.390 | P=0.023 | P=0.018 |
|  | cortex | P=0.313 | P=0.002 | P<0.001 |
| Glutamate | hippocampus | P<0.001 | P=0.029 | P<0.001 |
|  | cortex | P=0.072 | P<0.001 | P=0.421 |
| Glutathione | hippocampus | P=0.016 | P=0.398 | P=0.296 |
|  | cortex | P=0.184 | P<0.001 | P=0.015 |
| Glycine | hippocampus | P=0.243 | P=0.481 | P=0.624 |
|  | cortex | P=0.542 | P<0.001 | P=0.002 |
| *myo*-inositol | hippocampus | P=0.026 | P=0.063 | P=0.242 |
|  | cortex | P=0.750 | P<0.001 | P=0.528 |
| Lactate | hippocampus | P<0.001 | P<0.001 | P<0.001 |
|  | cortex | P=0.367 | P=0.002 | P=0.329 |
| NAA | hippocampus | P<0.001 | P<0.001 | P<0.001 |
|  | cortex | P=0.033 | P=0.034 | P=0.113 |
| Taurine | hippocampus | P<0.001 | P<0.001 | P<0.001 |
|  | cortex | P=0.989 | P<0.001 | P=0.281 |
| Ascorbate | hippocampus | P=0.084 | P=0.001 | P=0.888 |
|  | cortex | P=0.989 | P<0.001 | P=0.281 |
| Glucose | hippocampus | P=0.484 | P<0.001 | P=0.069 |
|  | cortex | P=0.680 | P=0.001 | P=0.011 |
| NAAG | hippocampus | P=0.042 | P=0.364 | P=0.151 |
|  | cortex | P=0.424 | P=0.573 | P=0.351 |
| PE | hippocampus | P=0.716 | P=0.138 | P=0.229 |
|  | cortex | P=0.602 | P<0.001 | P=0.474 |
| Total choline | hippocampus | P=0.002 | P<0.001 | P<0.001 |
|  | cortex | P=0.165 | P<0.001 | P=0.632 |
| Total creatine | hippocampus | P<0.001 | P<0.001 | P<0.001 |
|  | cortex | P=0.661 | P=0.018 | P=0.768 |
| PCr/Cr | hippocampus | P=0.033 | P=0.274 | P=0.932 |
|  | cortex | P=0.836 | P<0.001 | P=0.560 |

**Figure 5.**
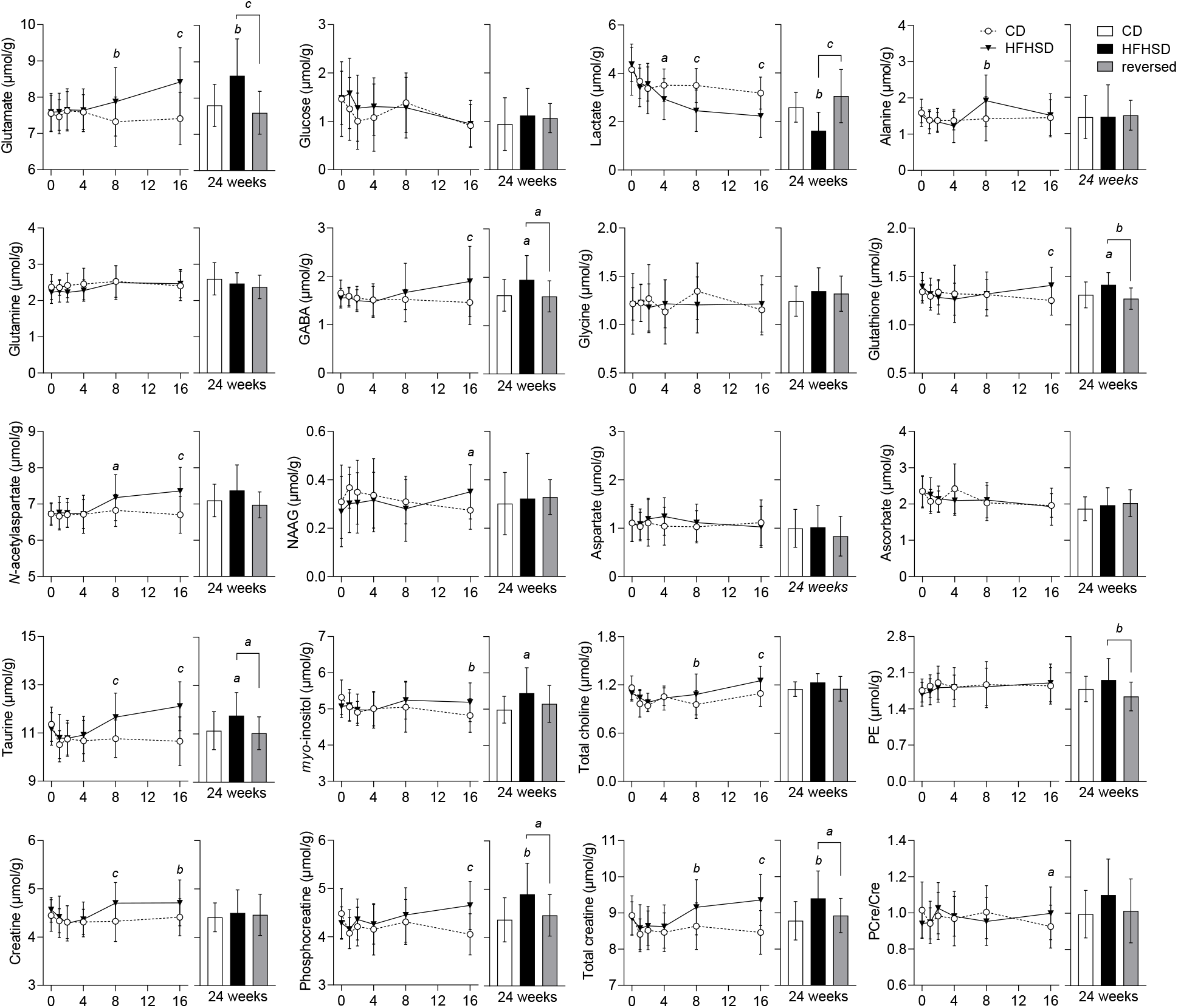
Alterations in metabolite levels triggered by HFHSD feeding in the hippocampus. Data is mean ± SD. Letters over data-points indicate significant differences relative to CD or as indicated (^*a*^ P<0.05, ^*b*^ P<0.01, ^*c*^ P<0.001) based on Fisher’s LSD post hoc comparison following presence of significant effects of diet or interaction between diet and time in ANOVA tests.

**Table 2.** ANOVA P-values for metabolite concentrations in the hippocampus and cortex at 24 weeks of the study.

| Metabolite | Brain area | ANOVA at 24 weeks |
| --- | --- | --- |
| Alanine | hippocampus | P=0.966 |
|  | cortex | P<0.001 |
| Aspartate | hippocampus | P=0.343 |
|  | cortex | P=0.580 |
| Creatine (Cr) | hippocampus | P=0.776 |
|  | cortex | P=0.003 |
| Phosphocreatine (PCr) | hippocampus | P=0.005 |
|  | cortex | P=0.903 |
| GABA | hippocampus | P=0.014 |
|  | cortex | P=0.781 |
| Glutamine | hippocampus | P=0.150 |
|  | cortex | P=0.051 |
| Glutamate | hippocampus | P<0.001 |
|  | cortex | P=0.069 |
| Glutathione | hippocampus | P=0.003 |
|  | cortex | P=0.002 |
| Glycine | hippocampus | P=0.205 |
|  | cortex | P=0.018 |
| *myo*-inositol | hippocampus | P=0.029 |
|  | cortex | P=0.138 |
| Lactate | hippocampus | P<0.001 |
|  | cortex | P=0.058 |
| NAA | hippocampus | P=0.064 |
|  | cortex | P=0.005 |
| Taurine | hippocampus | P=0.017 |
|  | cortex | P=0.542 |
| Ascorbate | hippocampus | P=0.433 |
|  | cortex | P=0.533 |
| Glucose | hippocampus | P=0.490 |
|  | cortex | P=0.644 |
| NAAG | hippocampus | P=0.819 |
|  | cortex | P=0.061 |
| PE | hippocampus | P=0.012 |
|  | cortex | P=0.938 |
| Total choline | hippocampus | P=0.063 |
|  | cortex | P=0.326 |
| Total creatine | hippocampus | P=0.005 |
|  | cortex | P=0.154 |
| PCr/Cr | hippocampus | P=0.122 |
|  | cortex | P=0.035 |

In general, the metabolite profile in the cortex (figure 6) was less impacted by HFHSD-feeding as compared to the hippocampus, and changes were generally in opposite direction. While cortical lactate levels remained unaltered, alanine concentrations were increased from 8 weeks of treatment in HFHSD-fed mice compared to controls (P<0.01). When compared to controls, we further observed a HFHSD-induced decrease of cortical *N*-acetylaspartate (P<0.05) from 8 weeks onwards and of glutamine (P<0.01) and glutathione (P<0.01) from 16 weeks onwards. Cortical glycine concentration showed a transient increase at 1 and 8 weeks of HFHSD feeding (P<0.05 *vs.* controls). While glycine levels were similar in HFHSD-fed mice and controls at 24 weeks of treatment, they were increased by diet reversal (P<0.05 *vs*. control and *vs*. HFHSD).

**Figure 6.**
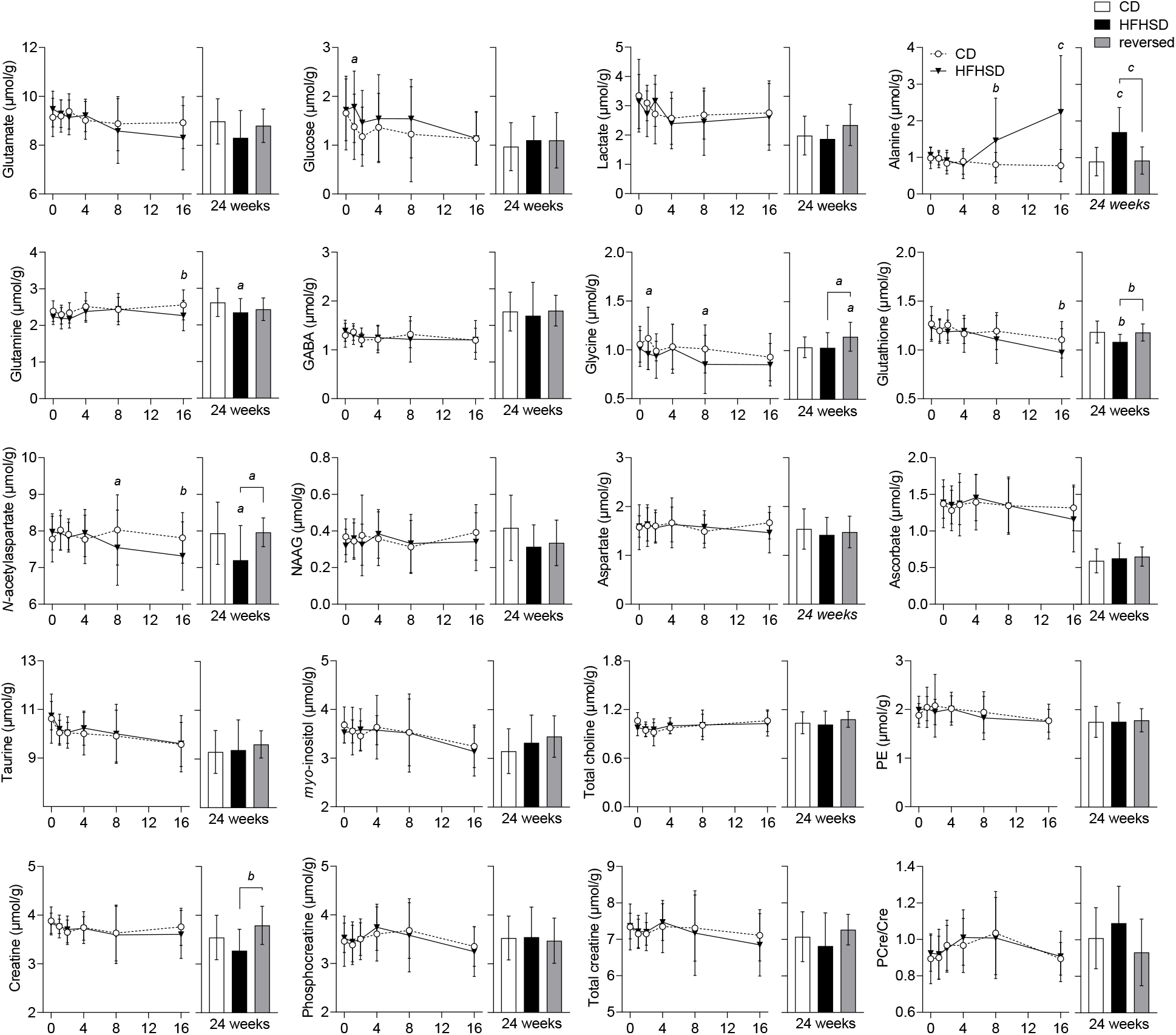
Alterations in metabolite levels triggered by HFHSD feeding in the cortex. Data is mean ± SD. Letters over data-points indicate significant differences relative to CD or as indicated (^*a*^ P<0.05, ^*b*^ P<0.01, ^*c*^ P<0.001) based on Fisher’s LSD post hoc comparison following presence of significant effects of diet or interaction between diet and time in ANOVA tests.

Brain glucose levels depend on glycemia and rates of brain glucose uptake and consumption (*e.g.* Duarte, Morgenthaler *et al.*, 2012). In line with increased glycemia, we observed a tendency of higher brain glucose levels in mice fed HFHSD, as compared to mice fed the control diet (table 1; figures 5–6).

We used principal component analysis (PCA) to provide a global depiction of brain metabolism. This analysis revealed a brain metabolic shift (relative to controls) in the hippocampus starting at 8 weeks of HFHSD feeding (figure 7A). The PCA conducted on MRS data collected during the first 16 weeks of treatment was then used to calculate the PCA scores from metabolite profiles acquired at 24 weeks. This analysis revealed a normalization of the hippocampal metabolite profile after diet reversal (Figure 7A). Results were less distinct for the cortex, in line with the less pronounced metabolic alterations observed in this region during the first 16 weeks of the dietary intervention (Figure 7B). Taurine and NAA were depicted by the PCA as the most influential metabolites in the effect of HFHSD on the brain metabolite profiles (figure 7C).

**Figure 7.**
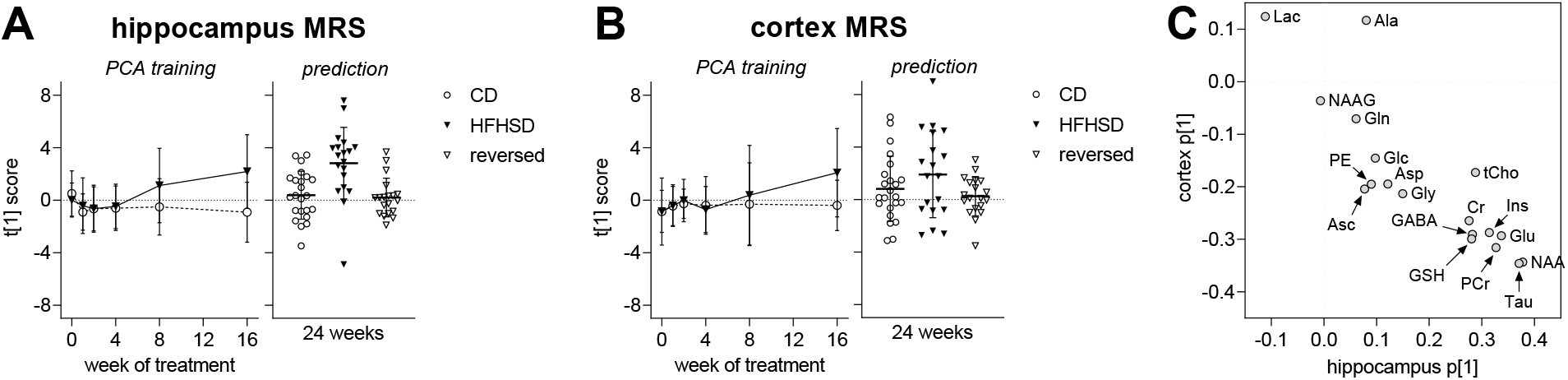
Alterations in the overall metabolite profile in hippocampus (A) and cortex (B) examined using principal component analysis (PCA). The PCA was calculated for the first 16 weeks. Trajectories are based on the scores along the first principal component (PC1), describing time-dependent alterations in the metabolite profile. Scores were then estimated from metabolite levels measured at 24 weeks of examination (prediction), revealing a normalization of the metabolite profile after diet reversal. Variation explained by the first principal component (PC1) was 29% in hippocampus and 36% in cortex. (C) Weights for each metabolite along PC1 for hippocampus and cortex. Data is individual data-points and mean±SD.

## Discussion

A number of studies suggest that obesogenic diets impact brain function (Garcia-Serrano & Duarte, 2020). However, it is hitherto unknown whether alterations in the brain are chronic or reversible by the introduction of a healthier diet. Hence, in the present study we tested the reversibility of HFHSD-elicited changes in brain metabolism and behavior. We found that metabolite profile alterations in the hippocampus and cortex develop gradually and are observable by MRS within a relatively short period of time, *i.e.* within 4 weeks after initiation of HFHSD feeding. Most importantly, brain metabolism and behavior largely normalized upon diet reversal, suggesting cellular adaptations to metabolic syndrome rather than major brain injury in HFHSD-fed mice. Moreover, despite of a more severe metabolic syndrome in HFHSD-fed male mice as compared to female mice, behavioral changes were similar, and both genders exhibited recovery potential upon diet reversal.

### Metabolic syndrome and brain function

The severity of the metabolic syndrome and brain dysfunction in mice subjected to obesogenic diets is known to depend on the amount and types of fat and sugar in the diet (Garcia-Serrano & Duarte, 2020; de Bem *et al.*, 2021). In this study we have implemented a model of hyper-caloric feeding in which additional calories originate mainly from saturated fat in a lard-based diet and from a sucrose solution provided *add libitum.* When compared to other diet-induced obesity models that have employed high-fat diets without added sugar on the same mouse strain, the present HFHSD model provides similar weight gain and glucose intolerance, but milder fasting hyperglycemia and hyperinsulinemia *(e.g.* Soares *et al.*, 2018; Lizarbe, Soares *et al.*, 2019). Altogether, this indicates a less severe metabolic syndrome in mice fed a HFHSD as compared to mice fed the more traditional HFD. Accordingly, the memory impairment in the Y-maze test was previously observed in mice under HFD (Lizarbe, Soares *et al.*, 2019) but not in the present study. On the other hand, object recognition tasks still depicted impaired memory performance in the HFHSD-fed mice. By analyzing free open-field exploration, we further found HFHSD-induced impairments in novel environment exploration and increased anxiety-like behavior (Prut & Belzung, 2003), as was also reported for mice exposed to HFD (Huber et al., 2021), and in line with a proposed obesity associated depression *(e.g.* Vagena *et al.*, 2019).

In contrast to most diet-induced obesity studies that have focused on male rodents, our study involved mice of both genders. Despite similar weight gain, female mice fed a HFHSD developed a metabolic phenotype that was less severe than in males. However, HFHSD feeding had a similar effect on behavior, that is, both males and females developed memory impairment and anxiety-like behavior. Interestingly, development of hyperinsulinemia was strongly sex-dependent, suggesting insulin levels to be a minor contributor to HFHSD-elicited brain dysfunction. However, the interaction between metabolic syndrome and the gender dimorphism of brain insulin sensitivity remains to be determined (Duarte *et al.*, 2018).

Several diabetes models show impaired memory performance that is thought to be linked to synaptic dysfunction rather than neuronal loss in the hippocampus (de Bem *et al.*, 2021). Moreover, obesity-independent impairments in hippocampal neurogenesis has been observed in T2D models, and proposed to contribute to poor performance in behavioral tasks (*e.g.* Lang *et al.*, 2009; Bracke *et al.*, 2019). Adult neurogenesis occurs mainly in the subventricular zone of the lateral ventricles and the subgranular zone of the dentate gyrus within the hippocampus, and has been linked to learning and memory, as well as behaviors related to stress, anxiety and depression (Snyder & Drew, 2020). Gender-specific modifications of hippocampal neurogenesis have been observed in mice exposed to HFD from 2 to 6 months of age (Robison *et al.*, 2020). The authors propose that deficits in cell proliferation within the dorsal but not ventral hippocampus, which was observed in females but not in males, could contribute to memory decline in females. In contrast, HFD feeding for 4 weeks led to impaired hippocampal neurogenesis in male but not female rats (Lindqvist et al., 2006). However, none of the studies reported hippocampal-dependent behavior alterations. Since factors other than neurogenesis contribute to adequate learning and memory performance, it is likely that cerebral homeostatic alterations triggered by HFHSD feeding lead to similar behavior impairments in male and female mice. Notably, brain dysfunction triggered by HFHSD feeding of mice of either gender was nearly fully recovered upon diet normalization. Recovery of memory impairments in male rats fed a HFD for 3 months after weaning were also partly recovered by diet normalization (Boitard *et al.*, 2016). Together with the absence of diabetes-induced neuronal death (de Bem *et al.*, 2021), this suggests that the above proposed synaptic dysfunction and reduced neurogenesis in obesity-associated metabolic syndrome can recover upon implementation of a healthy diet.

### Alterations in brain metabolism

Mice exposed to a HFHSD showed substantial alterations in the metabolite profile of both hippocampus and cortex. Furthermore, we now established that dimensionality reduction tools (PCA analysis) allowed the prediction of brain metabolite shifts, suggesting that the observed alterations in brain metabolism caused by HFHSD feeding are robust.

The hippocampus appeared to be more sensitive to HFHSD than the cortex, as also reported for HFD feeding (Lizarbe, Soares *et al.*, 2019). Reduced levels of lactate in the hippocampus of HFHSD fed mice from 4 weeks onward implies alterations in central energy metabolism. While the reduction in hippocampal lactate levels was sustained until the end of the study, we also observed a transient increase in alanine levels at 8 weeks of HFHSD feeding. In the cortex, alanine was increased after 4 weeks of HFHSD feeding in the absence of lactate changes. We have previously observed a reduction in the phosphocreatine-to-creatine ratio in mice after 6 months of feeding with a fat-enriched diet, when compared to mice fed a diet low in fat (Lizarbe, Soares *et al.*, 2019). These alterations were not observed in the present study, which is in line with the stronger impact of a HFD on brain energy metabolism, as compared with a diet that also include a high sugar content.

Total creatine content in the hippocampus was increased in mice fed HFHSD, relative to low-fat diet fed controls. This mimics previous observations in HFD fed mice (Lizarbe, Soares *et al.*, 2019), and suggests osmolarity dysregulation. Osmolarity control in the brain is considered to occur via the concentrations of not only creatine but also taurine, *myo*-inositol and *N*-acetylaspartate (discussed in Duarte, Lei *et al.*, 2012), which were found increased in the hippocampus of insulin resistant rats (Duarte *et al.*, 2019) and HFD-fed mice (Lizarbe, Soares *et al.*, 2019), relative to the respective controls. Indeed, levels of these four osmolytes were also increased in the hippocampus of HFHSD fed mice, and recovered upon diet reversal.

Brain taurine acts as an agonist of receptors involved in GABAergic and glycinergic neurotransmission (Albrecht & Schousboe, 2005), and plays and important role in preserving mitochondrial function and preventing oxidative damage (Hansen *et al.*, 2010). In addition to acting as organic osmolyte, increased taurine in the hippocampus of HFHSD-fed mice might thus be a beneficial adaptation, and act as compensatory mechanism for the loss of inhibitory tone and against mitochondrial stress. This beneficial adaptation was not observed in the cortex.

HFHSD-fed mice showed glutathione levels increased in the hippocampus but decreased in the cortex, relative to controls. Similar observations were reported in HFD-fed mice (Lizarbe, Soares *et al.*, 2019). It should be noted that the levels of glutathione measured by MRS include both reduced and oxidized forms, which are not distinguishable. Although a decreased ratio of reduced-to-oxidised glutathione has been reported as a consequence of short-term HFD feeding (Alzoubi *et al.*, 2018), brain levels of oxidized and reduced glutathione, and their ratio, have been shown to be unaffected by long-term HFD feeding (Li *et al.*, 2013). It is likely that a total glutathione increase in the hippocampus might reflect a compensatory mechanism to oxidative stress upon chronic HFHSD feeding. Interestingly, impaired cellular redox regulation in the brain has been proposed to compromise oligodendrocyte proliferation (consistent with increased levels of *N*-acetylaspartate, discussed below), which has implications on the maintenance of myelin sheets around axons, and thus brain connectivity (see Corcoba *et al.*, 2015, and references there in).

*N*-acetylaspartate is synthesized from neuronal mitochondrial acetyl-CoA and aspartate (Baslow, 2003), and its de-acetylation occurs in oligodendrocytes (Kirmani *et al.*, 2002). Therefore, reduced levels of *N*-acetylaspartate have been associated to neuronal injury, and its accumulation might represent oligodendrocyte dysfunction and impaired myelination, such as in Canavans disease (discussed in Duarte, Lei *et al.*, 2012). In the cortex, *N*-acetylaspartate levels were reduced by HFHSD feeding, indicating neuronal dysfunction. In the hippocampus, *N*-acetylaspartate accumulation could result from reduced aspartoacylase activity in oligodendrocytes, since high-fat feeding was suggested to impact myelination by promoting the loss of oligodendrocyte progenitor cells and mature oligodendrocytes (Yoon *et al.*, 2016).

We also found a HFHSD-induced consecutive increase of hippocampal concentrations of the main neurotransmitters glutamate and GABA. This suggests a differential impact of HFHSD feeding on glutamatergic and GABAergic systems. A tendency of increased levels of glutamate, but not GABA, has been reported in previous studies in which mice were fed a HFD for either 2 or 6 months (Liu et al., 2017; Lizarbe, Soares *et al.*, 2019). Liu *et al.* (2017) reported a HFD-induced increase in the rate of GABA synthesis from glucose without GABA accumulation in the brain. An earlier study on Zucker diabetic rats by Sickmann *et al.* (2010) suggested that diabetes has a differential impact on metabolic compartments that produce glutamate and GABA in the hippocampus, but not in the cortex or cerebellum. While T2D has been proposed to affect mainly glutamatergic synapses in NONcNZO/LtJ mice (Duarte, Agostinho *et al.*, 2012), the presynaptic vesicular carriers vGLUT1 and vGAT that are necessary for the function of both neurotransmission systems were affected by long-term HFD feeding (Lizarbe, Soares *et al.*, 2019). A direct link between glutamate and GABA levels and neurotransmission remains to be determined.

A transient increase in levels of *N*-acetylaspartylglutamate (NAAG) during HFHSD feeding was observed in the hippocampus. Increased NAAG concentrations were also observed in HFD-fed rats (Raider et al., 2016) and mice (Lizarbe, Soares *et al.*, 2019). NAAG is synthetized via ATP-dependent condensation of *N*-acetylaspartate and glutamate in neurons, and then packed into synaptic vesicles of presynaptic terminals, including those of pyramidal neurons in the cortex and limbic system. After being co-released with glutamate, NAAG acts as an antagonist of the glycine site of NMDA receptors, and interacts with the metabotropic type II glutamate receptor mGluR3 inhibiting the release of neurotransmitters, such as glutamate, GABA and glycine (Duarte & Xin, 2018, and referenced therein). Increased NAAG levels in the hippocampus of HFHSD-fed mice would thus reduce glutamatergic neurotransmission by both inhibiting the activity of NMDA receptors and augmenting the negative feedback of glutamate release through its action as an agonist of presynaptic mGluR3 receptors. High NAAG concentration might be a protective strategy, preventing toxic effects caused by excess of glutamate. On the other hand, sufficient glutamatergic activity is necessary for proper hippocampal function, and excessive NAAG might impair memory performance. Indeed, it was shown that both exogenously applied NAAG and inhibition of the NAAG-degrading enzyme glutamatecarboxypeptidase II abolish NMDA receptor-dependent long-term potentiation in the hippocampus, which is considered an *in vitro* form of memory (Bergeron *et al.*, 2007). Furthermore, by acting on astrocytic mGluR3, NAAG can also contribute in neurovascular coupling by dampening cerebral blood flow (Baslow *et al.*, 2005). While astrocytes of the hippocampus show high expression of mGluR3, its activity is proposed to be dependent on signaling mediated by other receptors (Bradlay & Challiss, 2012). Of particular interest is the finding that antagonism of the adenosine A2A receptor blunts mGluR3-mediated cAMP production (Moldrich *et al.*, 2002). Increased density of A2A receptors has been found in synaptic and mainly extra-synaptic membranes from the hippocampus in both obese and non-obese diabetes models (Duarte *et al.*, 2006; Duarte, Agostinho *et al.*, 2012; Duarte *et al.*, 2019). Therefore, one might speculate that increased NAAG levels together with an increased A2A receptor tone in astrocytes causes a dysfunctional regulation of astrocyte-mediated neurovascular coupling. In support of this, neurovascular dysfunction has been reported after 16, but not 8 weeks, of HFD feeding (Huber *et al.*, 2021).

The content of total choline was higher in hippocampus, but not cortex, of HFHSD-fed mice than controls, as has also been observed in HFD-fed mice (Lizarbe, Soares *et al.*, 2019). The main contributors to the choline peaks in MRS are the water-soluble glycerophosphorylcholine and phosphorylcholine, which are involved in membrane lipid metabolism, myelination processes and immune responses (Duarte, Lei *et al.*, 2012). Oxidized choline-containing phospholipids produced by oxidative stress have been proposed to contribute to neurodegeneration (Volinsky & Kinnunen, 2013). Recently, it was also proposed that oxidized choline containing lipids under normal conditions are cleared by microglia, to prevent neuronal injury (Dong *et al.*, 2021). Susceptibility to neuronal dysfunction might occur in HFHSD exposure, but taurine and GSH in the hippocampus might prevent oxidation of choline-containing lipids.

### Limitations

Others have reported behavior alterations after only one week of exposure to HFD in mice (McLean *et al.*, 2018). Against our expectations, we were unable to detect brain dysfunction by analyzing working spatial memory in the Y-maze, which was the behavior test of choice for longitudinal assessments after a pilot experiment using a HFD alone (data not shown). Therefore, the present study does not allow us to directly determine whether metabolic alterations precede neurocognitive dysfunction. We have observed the first sustained metabolic alteration at 4 weeks of HFHSD exposure (reduced lactate in hippocampus), but the Y-maze test revealed no signs of memory impairment at any time point.

Other limitation of the present study is that MRS metabolomics *in vivo* does not determine fluxes. Hence, it is unclear whether alterations in metabolite levels reflect metabolic flux modifications. However, the study still pinpoints metabolic pathways that are affected by HFHSD, which then can be more carefully investigated in future studies.

## Conclusion

Our study revealed robust and systematic alterations in the cortical and hippocampal metabolite profile in HFHSD-fed mice. Metabolite profile shifts indicate *e.g.* alterations in energy metabolism, adaptations to counteract osmolarity imbalances and mitochondrial stress, and cortical neurodegeneration. Importantly, even though these changes occurred within weeks after initiation of obesogenic diet feeding, we demonstrate an almost complete reversal of the observed phenotype after reinstatement of a healthier diet. Hence, HFHSD-elicited alterations in brain physiology show plasticity, which suggests adaptation rather than permanent structural damage of brain tissue in response to an unhealthy diet.

## Acknowledgements

The Lund University Bioimaging Centre is acknowledged for providing experimental resources. The authors are grateful Sara Larsson for experimental support and analysis of hormone levels, to Cecilia Skoug for valuable discussions regarding experimental planning, and to René In’t Zandt and Michael Gottschalk for MRS technical support.

## Author contribution statement

JMND designed the study and analyzed data. AMGS and AAM performed experiments and analyzed data. PS analyzed data. All authors wrote and revised the manuscript.

## Abbreviations

CD: control diet
GPC: glycerophosphorylcholine
GTT: glucose tolerance test
HFD: high-fat diet
HFHSD: high-fat and high-sucrose diet
LSD: least significant difference
MCI: mild cognitive impairment
MRS: magnetic resonance spectroscopy
NAAG: *N*-acetylaspartatylglutamate
NLR: novel location recognition
NOR: novel object recognition
PET: positron emission tomography
PCA: principal component analysis
PCho: phosphorylcholine
PE: phosphorylethanolamine
RD: reversed diet
T2D: type 2 diabetes
VOI: volume of interest.

